# Melatonin enhances peanut productivity by enriching root-associated nitrogen fixing bacteria

**DOI:** 10.1101/2025.08.11.669620

**Authors:** Ali Muhammad, Xiangjun Kong, Lijie Li, Muhammad Hafeez Ullah Khan, Peipei Jia, Chen Miao, Zhiyong Zhang

## Abstract

Melatonin, a pleiotropic phytohormone, is widely recognized as a promising bio-stimulant, yet its integrative effects on root development, yield gain, and microbiome assembly in legumes remain underexplored. In this study, we investigated the effects of melatonin seed treatment across three peanut genotypes, focusing on plant productivity, and the composition and structure of bacterial communities in root, rhizosphere, and bulk soil compartments. Melatonin treatment substantially improved root biomass, nodulation, nitrogen balance index, and yield-related traits, with the highest response observed in the genotype Kainong 308. Amplicon sequencing revealed that melatonin induced distinct genotype and compartment specific shifts in bacterial community composition, with the root bacteria showing the increased remodeling, including a 45.9% increase in unique amplicon sequence variants (ASVs). Melatonin selectively enriched key Proteobacteria taxa such as *Rhizobium*, *Sphingomonas*, and *Enterobacter hormaechei*, known for their plant-growth promoting and biocontrol capabilities. Co-occurrence network analysis indicated that melatonin-treated roots harbored more complex bacterial networks, and module #4 dominated by melatonin-induced Proteobacteria was strongly correlated with most of the plant traits. Collectively these findings highlight melatonin dual role as a bio-stimulant and microbiome modulator, promoting a functionally enriched and responsive bacteria that supports enhanced plant performance. This study provides novel insights into the melatonin-mediated coordination of plant performance and bacterial assembly, offering a foundation for microbiome-informed crop improvement strategies.

## Introduction

Crop production has traditionally focused on high-input conditions, neglecting the role of plant-associated microbial communities (Quiza *et al*., 2023; Xing and Wang, 2024). This has led to the view that modern genotypes have lost their potential to interact with beneficial microbial communities when grown under low-input conditions (Emmett *et al*., 2018; Quiza *et al*., 2021). However, plant associated microbes can have positive effects on plant performance, including nutrition acquisition, pathogen inhibition, growth promotion, and stress resilience, their absence can greatly hinder efforts toward sustainable agriculture. It is important to consider the impact of environmental conditions (Li *et al*., 2020), plant growth stage (Chaparro *et al*., 2014; Moroenyane *et al*., 2021a), plant compartment (Agoussar *et al*., 2021; Moroenyane *et al*., 2021a), and their interactions (Moroenyane *et al*., 2021c) on plant-associated microbial communities. Plant compartment, particularly the rhizosphere (the soil surrounding the roots), has been identified as a significant factor influencing the diversity of plant-associated microbial communities (Coleman-Derr *et al*., 2016; Moroenyane *et al*., 2021a). The rhizosphere effect refers to the differences observed between microbial communities in the rhizosphere and those in the bulk soil (Lv *et al*., 2023; Sun *et al*., 2023). Additionally, microbial communities in the roots, rhizosphere, and aboveground parts of plants are distinct (Bai *et al*., 2015; Moroenyane *et al*., 2021a), with some similarities found in seed microbial communities (Moroenyane *et al*., 2021b).

Plant-microbe crosstalk and the role of PGPR in the rhizosphere has been explored (Bhat *et al*., 2020; Srivastava *et al*., 2024), limited research has been done on the rhizosphere of peanuts (*Arachis hypogea*. L) (Dai *et al*., 2019; Haldar and Sengupta, 2015). Peanuts are economically important legumes that establish a symbiotic association with rhizobia through root exudates, which is important for their overall yield. In addition to root exuded chemicals, there is a growing interest in the effects of exogenous treatment with various substances of both natural and synthetic origin, such as phytohormones, osmolytes, and gaseous molecules (Bargunam *et al*., 2025; Mukherjee and Corpas, 2020; Raza *et al*., 2022). These substances promote seedling emergence and plant growth, reduce plant growth periods, and enhance productivity (Hussain *et al*., 2024). Among these substances melatonin (MT) a pleiotropic biomolecule is recognized as a key regulator (Shahani *et al*., 2023). MT is derived from serotonin (5-hydroxytryptamine), is widely distributed in plants and animals associated with numerous biological functions. These functions include seed germination, root growth, flowering, leaf senescence, reproduction, fruit ripening, circadian rhythm regulation, and plant adaptation to varying environments (Ahammed *et al*., 2020; Akula and Mukherjee, 2020). Seed treatment with melatonin offers additional benefits by enhancing photosynthetic activity and promoting growth (Shaheen *et al*., 2024; Zhang *et al*., 2021). Melatonin growth-promoting effects are particularly pronounced under stressful condition affecting plant development, as in the case of *Phaseolus vulgaris* (Alinia *et al*., 2022) and maize (Xu *et al*., 2024) subjected to salt stress, peanuts and cotton (de Camargo Santos *et al*., 2024; Zhu *et al*., 2023) under drought stress, and *Malus* grown in nutrient-deficient conditions (Du *et al*., 2022). Recently, Qiao *et al*. (2019) revealed melatonin activity in nitrogen uptake and metabolism-related enzymes synthesis promoting winter wheat growth under nitrogen deficient conditions. These findings endorse melatonin role in facilitating nutrient uptake and metabolism, promoting plant growth and development during stressful conditions. In line with these findings, earlier work from our research group demonstrated that melatonin-treated plants achieved higher peanut yield by maintaining more stable carbon and nitrogen metabolism compared to control ones (Li *et al*., 2024). Based on these findings, we hypothesized that seed treatment with melatonin, favors the highest abundance of symbiotic microorganisms associated with plants. To test this, we investigated genotype-specific effects of melatonin on root development and yield associated traits, alongside shifts in bacterial community composition and diversity across root, rhizosphere, and bulk soil compartments. This study provides a potential novel pathway for communication between microbial symbionts and their host plants mediated by melatonin.

## Materials and methods

### Experimental design and plant material

The field trial was conducted from the mid-May to the end of September 2023 at the Yanjin experimental field of the Henan Institute of Science and Technology in Xinxiang City, Henan Province, China (114°12′60″E, 35°21′43″N), a region characterized by a warm temperate continental monsoon climate with an average annual precipitation of 653.20 mm (Qin *et al*., 2023). The experiment was conducted in a randomized complete block design with three replications. Peanut varieties locally grown, namely Xinbaihua 16, Xinbaihua 21, and Kainong 308 were selected for the experiment. The soil texture was sandy loam. To investigate the effect of melatonin on plant performance and bacterial abundance, seeds were treated with 0.5µM of melatonin (MT), while untreated seeds served as the control (CK). In order to characterize the bacterial community, samples will be obtained from the root surface, rhizosphere soil, and bulk soil (in the middle of two rows) at maturity.

### Plant growth and yield parameters

Ten representative plant samples were collected from each treatment at maturity. Plant growth parameters including root related traits such as total root length, taproot length, belowground biomass, lateral root number, number of nodules, and root surface area. Yield contributing traits were also measured including 100-pod weight, number of pods per plant, 100-grain weight and total yield. For aboveground biomass measurement, randomly selected plants were carefully uprooted and weighted for total biomass. Chlorophyll florescence was evaluated on a fully expanded dark acclimated (for ≥ 20 min) leaves, using the handy PEA fluorimeter (Hansatech, King’s Lynn, UK) with a saturating light pulse of 3500 µmole m^−2^s^−1^.

### Plant and soil sampling

Plants and soil samples were collected at maturity using the five-point diagonal sampling method. The bulk soil was collected from a depth of 5-20 cm between two rows of crops, ensuring that no roots were included in the soil. For collecting rhizosphere soil, the roots were gently brushed to remove the loosely attached soil, leaving the roots with very little soil adhering to them. The roots were cut off from the shoot with sterile scissors and then placed with the adhering soil in 50 ml sterile centrifuge tube containing 40 ml PBS buffer. The tubes were then centrifuged at a high speed to remove the root surface soil, after which the roots were removed from the PBS solution. The PBS containing the root surface soil was centrifuged at 2000g for 5 min, after which the supernatant was discarded and the rhizosphere soil collected. All the soil samples and roots were immediately stored at −80 °C for high-throughput sequencing. Three biological replicates were performed for each treatment and a total of 54 samples were collected (three varieties × two treatments × three compartments × three replicates). For soil chemical analysis including nitrate nitrogen, ammonium nitrogen, and phosphorus, the soil loosely attached to the roots, was collected in a sterile container. The soil pH and electrical conductivity (EC) were determined using a soil-to-water ratio of 1:2.5 with a pH meter (Mettler Toledo, Zurich, Switzerland) and a conductivity meter (INESA, Shanghai, China).

### DNA extraction and 16S rRNA gene amplification

The soil microbial DNA was extracted with the TIANamp Soil DNA Kit using TGuide S96 paramagnetic particle method, following the manufacture instructions (Tiangen, Beijing, China). The concentration and quality of the DNA were assessed using a Nanodrop 2000 spectrophotometer (Thermo Scientific, Waltham, USA) and 1% agarose gels. The V3-V4 region of the bacterial 16S rRNA gene was amplified using the forward primer 338F (5’-ACTCCTACGGGAGGCAGCA-3’) and the reverse primer 806R (5’-GGACTACHVGGGTWTCTAAT-3’). PCR amplification was carried out using TransStart FastPfu Fly DNA Polymerase kit (TransGen Biotech Beijing, China) with the following conditions: denaturation at 95°C for 5 min, 25 cycles of 30s at 95°C, annealing at 50°C for 30s, elongation at 72°C for 40s, and final extension at 72°C for 5 min. The resultant PCR products were assessed by agarose gel electrophoresis and purified with the E.Z.N.A. cycle pure kit (Omega, USA). Finally, the libraries were sequenced on an Illumina NovaSeq 6000 platform (Biomarker, Beijing, China).

### Bioinformatic analysis

After sequencing the raw reads were primarily filtered by Trimmomatic v.0.33 (Bolger et al.,2014). The primer sequences were identified and denoised with Cutadapt v.1.9.1 (Callahan *et al*., 2016). In the present study, a total of 3,888,705 high quality reads were obtained. Sample rarefaction curves revealed asymptotic trend, which ensured that the sequencing depth was sufficient for subsequent bacterial diversity analysis (**Fig. S1**). Sequencing data were processed with the DADA2 pipeline to denoise reads, and generate high resolution ASVs (Callahan *et al*., 2016). ASVs with an abundance of <0.005% were removed, resulting in the identification of 56,650 ASVs (Bokulich *et al*., 2013). Taxonomic annotation of the representative sequences was performed using the Bayesian classifier with SILVA as the reference database (Quast *et al*., 2012). QIIME2 was utilized to determine the abundance of each specie in the samples (Bolyen *et al*., 2019).

### Statistical analysis

The effect of melatonin on soil properties and peanut growth parameters was tested by one-way analysis of variance (LSD test, P< 0.05) using SPSS v.22.0, while, other statistical analyses were performed using R Studio. The principal coordinate analysis (PCoA) was performed using the “vegdist” function in the “vegan” package, based on the Bray-Curtis dissimilarity matrices (Oksanen, 2015). The effects of genotype, compartment, and melatonin treatment on the community composition was tested by permutational multivariate analysis of variance (PERMANOVA) using the “adonis2” function of the “vegan” package. The bacterial species distribution histogram was calculated and visualized using the “tidyverse” and “ggsci” packages. Co-occurrence network was constructed based on the ASVs with a relative abundance > 0.01% and present in more than half of the samples (Liu *et al*., 2023). The relative abundance of each cluster was calculated by averaging the standardized (z-score) relative abundances of their corresponding ASVs (Fan *et al*., 2021). The functional potential of bacterial community was explored using PIRCUSt2 (Gao *et al*., 2024).

## Results

### Effects of melatonin on root growth and yield associated traits

Melatonin application had significant effects on root parameters and yield associated traits across the three peanut genotypes (XBH16, XBH21, and KN308). Nodulation was considerably enhanced by melatonin in all genotypes, with the number of nodules ranged from 167.33 to 204 in XBH16, and from 117.5 to 151.5 in KN308 (**Fig. 1a**). Taproot length also enhanced remarkably under melatonin, attaining 17.62 cm and 14.8 cm for XBH16 and KN308, respectively, compared to 14.66 cm and 12.01 cm of their corresponding control (**Fig. 1b**). Root biomass showed a substantial increase, especially in KN308, where root weight increased from 1.15 g in CK to 1.73 g in MT (**Fig. 1c**). Regarding yield components, 100-grain weight increased under melatonin from 80.76 to 88.17 g in XBH16, and from 75.28 to 81.97 g in KN308 (**Fig. 1d**). Similarly, 100-pod weight reveled improvement among genotypes, with KN308 showing an increase from 171.71 to 182.14 g (**Fig. 1e**). Finally, yield was significantly enhanced by melatonin in XBH16 (149.46 to 175.21 g/m^2^) and KN308 (151.53 to 187.04 g/m^2^), with KN308 exhibiting the highest yield among all treatments (**Fig. 1f**). These results were supported by significant genotype, treatment, and genotype × treatment interaction effects for important traits such as nodules, root weight, and yield (**Table 1**). Additionally, melatonin treatment led to improvements in total root length, aboveground biomass, nitrogen balance index, and chlorophyll content, although the extent of response varied among genotypes (**Table 1**).

**Fig. 1.**
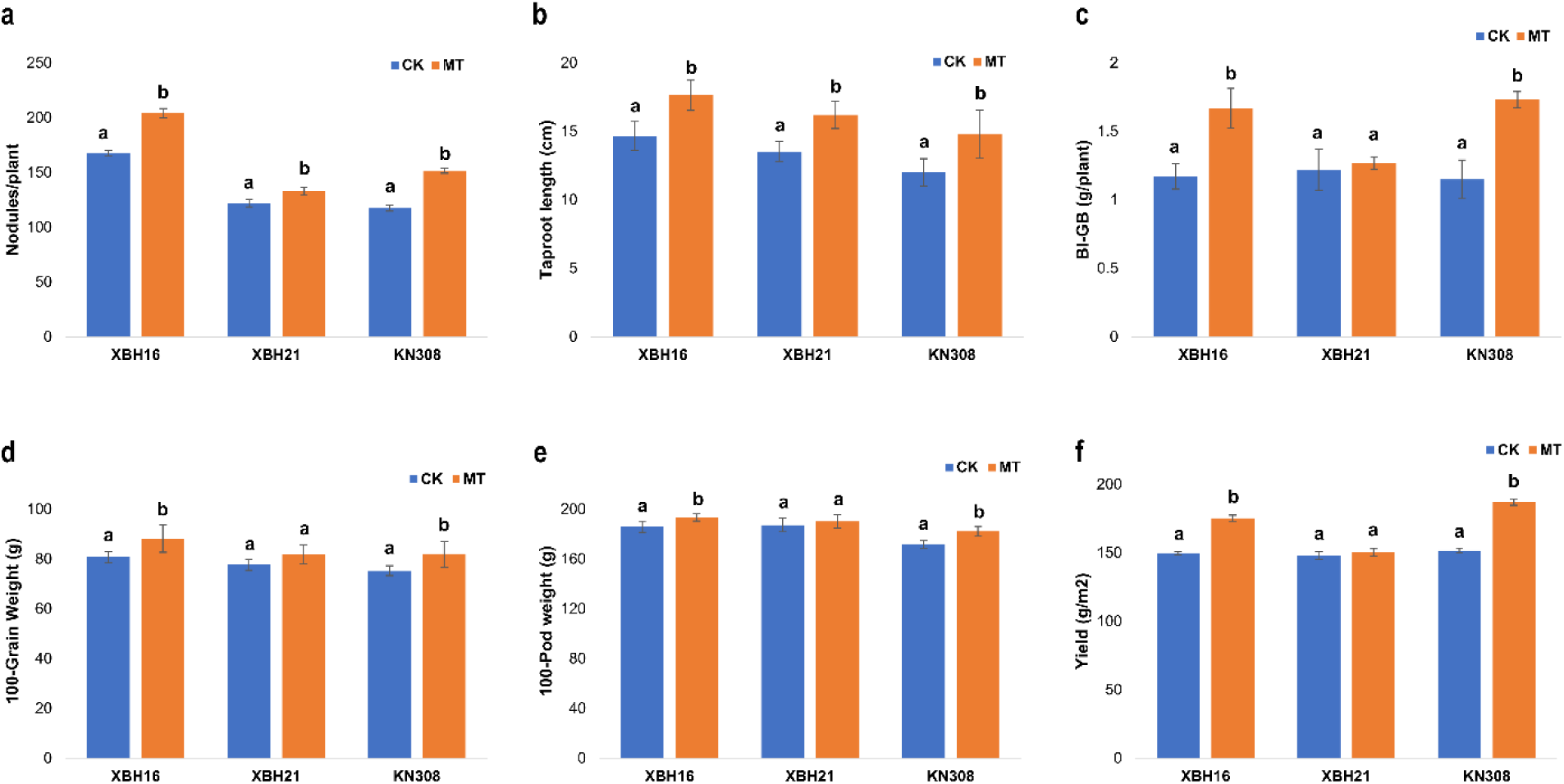
Effects of melatonin treatment on root and yield-related traits in three peanut genotypes (XBH16, XHB21, and KN308). (a) Number of nodules per plant, (b) Taproot length (cm), (c) Belowground biomass (g/plant), (d) 100-grain weight (g), (e) 100-pod weight (g), and (f) Yield (g/m^2^) under control (CK) and melatonin (MT). Data are presented as mean values. Different lowercase letters above bars show significant differences (*p* < 0.05) between CK and MT. Error bars indicate standard error. XBH16, xinbaihua 16; XBH21, xinbaihua 21; KN308, kainong 308.

**Table 1.** Summary of physiological, root, and yield-related traits in three peanut genotypes (XBH16, XBH21, and KN308) under control (CK) and melatonin (MT) treatments.

| Genotype<br>Treatment | XBH16 |  | XBH21 |  | KN308 |  | P-Value |  |  |
| --- | --- | --- | --- | --- | --- | --- | --- | --- | --- |
|  | CK | MT | CK | MT | CK | MT | Geno | Treat | Geno*Treat |
| **pH** | 7.72 ± 0.13 | 7.70 ± 0.10 | 7.46 ± 0.19 | 7.42 ± 0.11 | 7.73 ± 0.17 | 7.72 ± 0.01 | 0.000 | 0.417 | 0.839 |
| **EC** | 7.09 ± 1.30 | 6.75 ± 1.50 | 6.77 ± 0.77 | 6.62 ± 0.58 | 7.35 ± 3.47 | 6.66 ± 1.47 | 0.749 | 0.269 | 0.808 |
| **NN** | 122.98 ± 5.01 | 164.52 ± 16.02 | 171.78 ± 9.67 | 261.53 ± 14.99 | 191.15 ± 8.30 | 269.57 ± 8.31 | 0.000 | 0.000 | 0.000 |
| **AN** | 12.91 ± 2.60 | 13.81 ± 3.67 | 11.41 ± 1.63 | 12.72 ± 3.08 | 9.47 ± 1.67 | 10.80 ± 2.76 | 0.001 | 0.038 | 0.928 |
| **P** | 168.68 ± 7.78 | 159.23 ± 15.49 | 203.78 ± 15.21 | 198.09 ± 11.63 | 160.46 ± 11.44 | 165.09 ± 12.85 | 0.000 | 0.171 | 0.084 |
| **Fv/Fm** | 0.79 ± 0.03 | 0.80 ± 0.05 | 0.79 ± 0.04 | 0.82 ± 0.07 | 0.72 ± 0.01 | 0.79 ± 0.08 | 0.002 | 0.003 | 0.077 |
| **NBI** | 46.42 ± 3.76 | 62.73 ± 9.46 | 41.93 ± 9.59 | 70.62 ± 6.16 | 49.86 ± 3.82 | 54.81 ± 9.75 | 0.121 | 0.000 | 0.000 |
| **Chl** | 31.24 ± 2.70 | 33.98 ± 4.09 | 29.50 ± 7.46 | 38.00 ± 5.75 | 27.10 ± 2.76 | 31.55 ± 2.21 | 0.000 | 0.06 | 0.882 |
| **RN** | 37.11 ± 2.9 | 38.06 ± 3.05 | 40.11 ± 3.76 | 41.83 ± 4.36 | 36.1 ± 3.45 | 37.33 ± 2.18 | 0.002 | 0.063 | 0.656 |
| **Nodules** | 167.33 ± 6.25 | 204.00 ± 10.83 | 122.00 ± 8.72 | 133.17 ± 8.45 | 117.50 ± 6.92 | 151.50 ± 5.39 | 0.000 | 0.000 | 0.000 |
| **TPL** | 14.66 ± 2.65 | 17.62 ± 2.73 | 13.53 ± 1.80 | 16.22 ± 2.52 | 12.01 ± 2.47 | 14.80 ± 4.38 | 0.005 | 0.000 | 0.978 |
| **RSA** | 25.18 ± 4.09 | 26.68 ± 1.83 | 23.80 ± 3.99 | 26.17 ± 1.59 | 21.47 ± 2.61 | 25.40 ± 3.62 | 0.015 | 0.001 | 0.277 |
| **TRL** | 576.29 ± 9.19 | 591.18 ± 8.97 | 616.93 ± 12.22 | 627.52 ± 11.55 | 491.69 ± 22.14 | 638.96 ± 3.97 | 0.000 | 0.000 | 0.000 |
| **Ab-Gb** | 12.79 ± 1.80 | 21.41 ± 5.53 | 14.03 ± 3.63 | 18.00 ± 3.03 | 20.55 ± 2.50 | 25.07 ± 6.31 | 0.000 | 0.000 | 0.063 |
| **Bl-GB** | 1.17 ± 0.23 | 1.67 ± 0.36 | 1.22 ± 0.37 | 1.27 ± 0.11 | 1.15 ± 0.35 | 1.73 ± 0.15 | 0.021 | 0.000 | 0.003 |
| **GW** | 80.76 ± 5.67 | 88.17 ± 13.58 | 77.7 ± 5.49 | 81.83 ± 9.25 | 75.28 ± 5 | 81.97 ± 12.76 | 0.044 | 0.004 | 0.734 |
| **PW** | 185.83 ± 10.73 | 193.34 ± 6.93 | 187.22 ± 13.42 | 190.17 ± 13.09 | 171.71 ± 7.52 | 182.14 ± 9.82 | 0.000 | 0.004 | 0.341 |
| **Yield (g/m<sup>2</sup>)** | 149.46 ± 1.21 | 175.21 ± 2.18 | 147.92 ± 2.56 | 150.27 ± 2.66 | 151.53 ± 1.73 | 187.04 ± 2.32 | 0.000 | 0.000 | 0.000 |

Data are presented as mean ± 95% confidence interval. Abbreviations: EC, electrical conductivity; NN, nitrate nitrogen; AN, ammonium nitrogen; P, Phosphorus; NBI, nitrogen balance index; Chl, chlorophyll content; RN, root number; TPL, taproot length (cm); RSA, root surface area (cm<sup>2</sup>); TRL, total root length (cm); Ab-GB, aboveground biomass (g); Bl-GB, belowground biomass (g); GW, 100-grain weight (g); PW, 100-pod weight (g).

### Melatonin-induced shifts in bacterial community composition across rhizocompartments

To know the influence of melatonin on the overall bacterial community composition, we conducted Principal coordinates analysis (PCoA), Permutational multivariate analysis of variance (PERMANOVA), and Linear discriminant analysis effect size (LEfSe)-based taxonomic profiling. The PCoA plot (**Fig. 2a**) revealed clear separation of bacterial communities among roots (R), rhizosphere soil (RS), and bulk soil (BS), with PC1 explaining 28.36% of the total variation. Notably, bacterial communities in the root compartment, clustered distinctly from those in RS and BS, revealing strong compartment-specific microbial structuring. Within each compartment, MT and CK samples indicated distinct clustering, especially in the root samples of XBH16 (R16) and KN308 (R308), suggesting genotype specific effects of melatonin on bacterial abundance. This was further supported by the PERMANOVA results (**Fig. 2b**), which confirmed significant differences between bacterial communities across treatments (R = 0.7036, *p* = 0.001). Moreover, bacterial composition varied more strongly across treatments than within individual treatment groups.

**Fig. 2.**
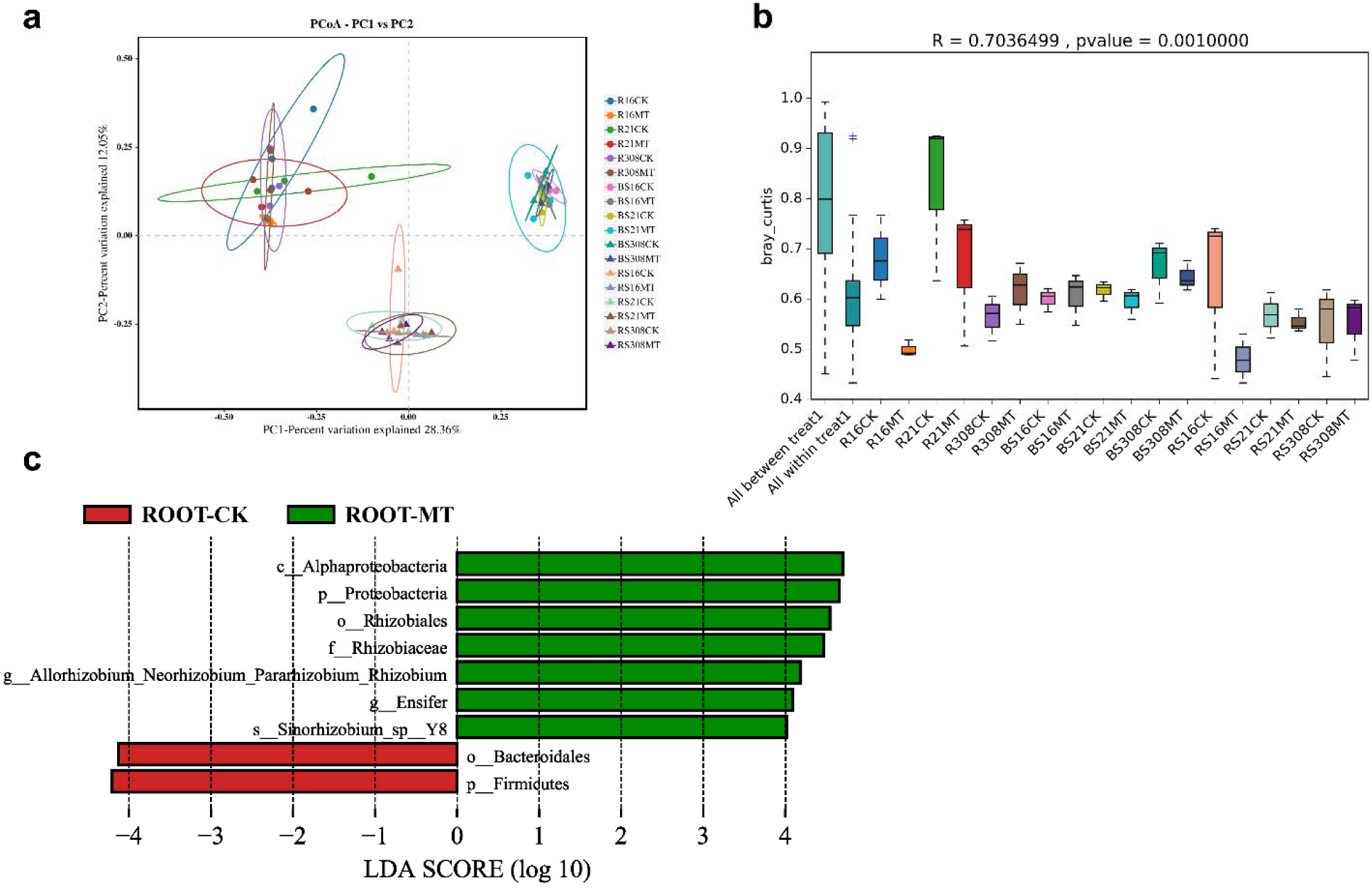
Clustering analysis of bacterial community composition in peanut rhizocompartments under melatonin (MT) and control (CK) treatments. (a) Principal Coordinates Analysis (PCoA) based on Bray-Curtis distance showing distinct clustering of bacterial communities across root (R), rhizosphere soil (RS), and bulk soil (BS) samples from three peanut varieties (XBH16, XBH21, and KN308) under control (CK) and melatonin (MT) treatments. Each point represents a sample, and ellipses indicate 95% confidence intervals for each group. (b) Permutational multivariate analysis of variance (PERMANOVA)-based boxplot illustrating Bray-Curtis dissimilarity within and between treatment groups. A significant separation was observed between treatments (R = 0.7036, P = 0.001), indicating that melatonin treatment significantly altered bacterial community composition. (c) Linear discriminant analysis effect size (LEfSe) revealed important bacterial taxa significantly enriched in root compartments under melatonin (green) and control (red) treatments.

LEfSe analysis revealed that MT treatment significantly enriched Proteobacteria-related taxa (*Rhizobium*, *Pararhizobium*, *Ensifer*) in the root compartments, while roots under CK were dominated by *Firmicutes* and *Bacteroidales* (**Fig. 2c**). These shifts show that melatonin selectively promotes the proliferation of beneficial bacterial taxa in the root compartment, potentially influencing plant-microbe interactions. A more detailed analysis across all rhizocompartments and genotypes further identified several important Proteobacteria, including *Allorhizobium* and *Enterobacter hormaechei*, along with members of Sphingomonadaceae significantly enriched under melatonin treatment. In contrast, CK samples indicated greater abundance of unclassified Chloroflexi and certain Actinobacteria (**Fig. S2**). These shifts were more obvious in the root and rhizosphere compartments, suggesting a compartment and genotype specific bacterial response to melatonin, likely contributing to improved nutrient acquisition, nodulation, and plant growth.

### Taxonomic composition of bacterial communities in response to melatonin

The bacterial composition was characterized based on the relative abundance profiles across five taxonomic levels (phylum, class, order, family and genus). At the phylum level, Proteobacteria (19.03-64.91%) consistently dominated across all three genotypes, followed by Acidobacteriota (4.17-26.95%), and Actinobacteriota (6.09-15.57%) (**Table S1**). Compartment-specific trends revealed that Proteobacteria were predominant in root and rhizosphere samples, whereas Acidobacterota were more abundant in bulk soil. Melatonin treatment influenced the composition of bacterial community in a genotype and compartment dependent manner. Generally, melatonin increased the relative abundance of most dominant phyla. However, a notable decline in the abundance of Bacteriodota and Firmicutes was observed under MT treatment, suggesting a potential inhibitory effect of melatonin on these bacterial taxa (**Fig. 3**).

**Fig. 3.**
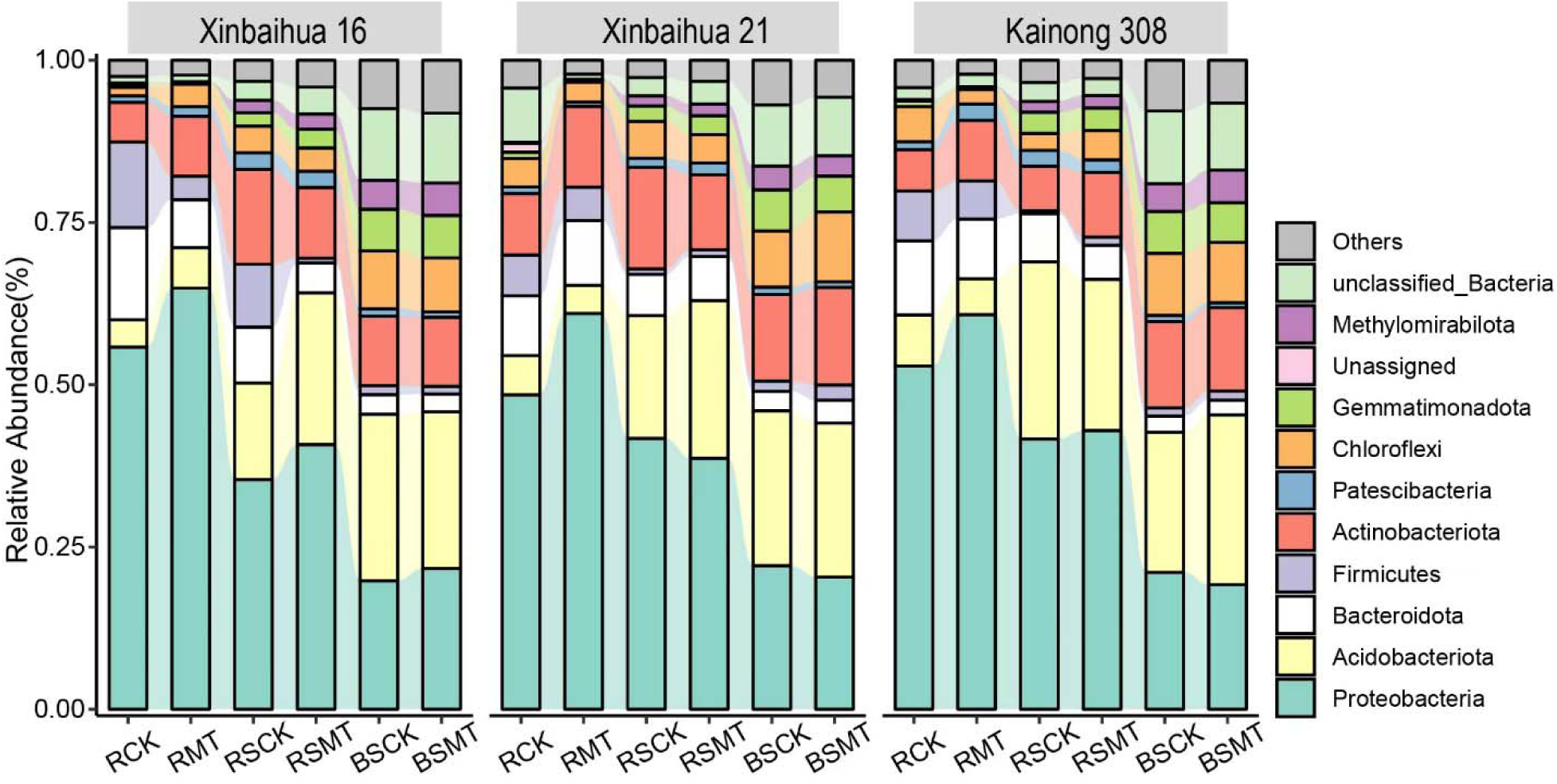
Relative abundance of bacterial communities at the phylum level across peanut genotypes and rhizocompartments under control and melatonin treatments. Stacked bar plots represent the relative abundance of dominant phyla in root (R), rhizosphere soil (RS), and bulk soil (BS) of three peanut genotypes, Xinbaihua 16, Xinbaihua 21, and Kainong 308, under control (CK) and melatonin (MT) treatments. “Unassigned” denotes sequences that could not be classified at the phylum level, while “Others” indicate low abundance classified phyla not individually labeled.

Further analysis of bacterial community composition at finer taxonomic resolution revealed clear melatonin-induced shifts across all genotypes and compartments (**Fig. S3**). At the class level, Alphaproteobacteria and Gammaproteobacteria indicated and obvious increase in relative abundance under MT treatment, especially in root and rhizosphere samples, suggesting a potential stimulatory effect of melatonin on these classes (**Fig. S3a**). At the order level, Vicinamibacterales and Rhizobiales dominated the community across samples (**Fig. S3b**). Vicinamibacteraceae emerged as the most abundant family across compartments, while sphingomonadaceae was particularly enriched in the root and rhizosphere samples (**Fig. S3c**). Genus-level characterization revealed an increased abundance of *Sphingomonas* in MT-treated root and rhizosphere compartments, suggesting a positive bacterial response to melatonin (**Fig. S3d**). Finally, some sequences remained unclassified, highlighting the unexplored bacterial diversity of peanut-associated soils and the need for further characterization.

### Bacterial community composition and trait associations across compartments

Melatonin application influenced the bacterial community structure in a compartment and genotypic specific manner. Petal plots (**Fig. 4a-c**) revealed considerable variation in OUT richness among, root, rhizosphere, and bulk soil samples. Within the root compartment, R21CK exhibited the highest number of unique ASVs (3595), while R308MT had the lowest (1987). Despite this variability, only 306 ASVs were consistently shared across all root samples, indicating strong selective pressures imposed by both genotype and treatment. The rhizosphere compartment displayed a moderately overlapping community, with 495 core ASVs, whereas bulk soil compartment maintained a relatively stable microbiota, sharing 539 ASVs across all genotypes and treatments. This observation was further supported by the Venn diagram (**Fig. 4d**), where the root compartment included the greatest proportion of unique ASVs (45.9%), compared to the rhizosphere (17.6%) and bulk soil (24.3%). However, only 1% (29 ASVs) were common across all compartments, showing pronounced ecological separation among compartments.

**Fig. 4.**
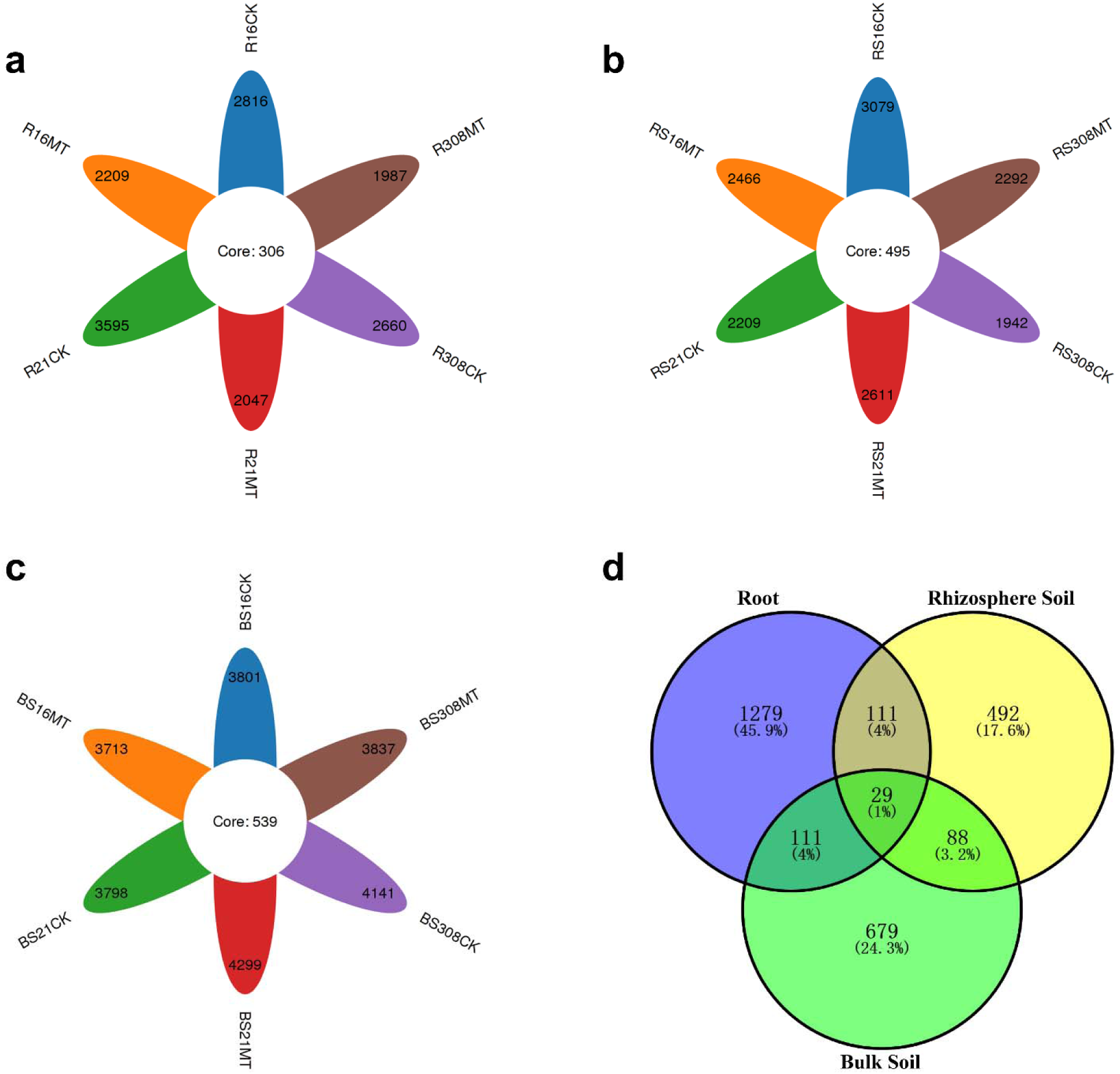
Comparative analysis of bacterial community composition across compartments and treatments in peanut. (a-c) Petal plots illustrate the number of shared and unique ASVs in root (R), rhizosphere soil (RS), and bulk soil (BS) under control (CK) and melatonin (MT) across three peanut genotypes (Xinbaihua 16, Xinbaihua 21, and Kainong 308). Each petal shows ASVs specific to a treatment-genotype combination, while the central region represents core ASVs shared within each compartment. (d) Venn diagram revealing ASV overlap across the three compartments, highlighting distinct bacterial niche separation between root, rhizosphere, and bulk soil environments.

Given the obvious compartment-specific differences in bacterial composition, we further explored how these communities relate to plant phenotypic and soil traits (**Fig. 5**). Among soil chemical parameters. Ammonium nitrogen showed positive correlation with most of plant traits, including NBI, chlorophyll content, root number, biomass accumulation, and yield. Key physiological traits, such as chlorophyll content, NBI, and Fv/Fm, as well as root-related parameters, were also strongly correlated with 100-pod weight, and yield. Mentel test analysis further confirmed that bacterial communities in the root compartment were more closely associated with plant traits than those in the rhizosphere and bulk soil. Together, these findings underscore the compartment-driven influence of melatonin on bacterial abundance, particularly within the root microbiome, which revealed the strongest response and most distinct association with plant performance.

**Fig. 5.**
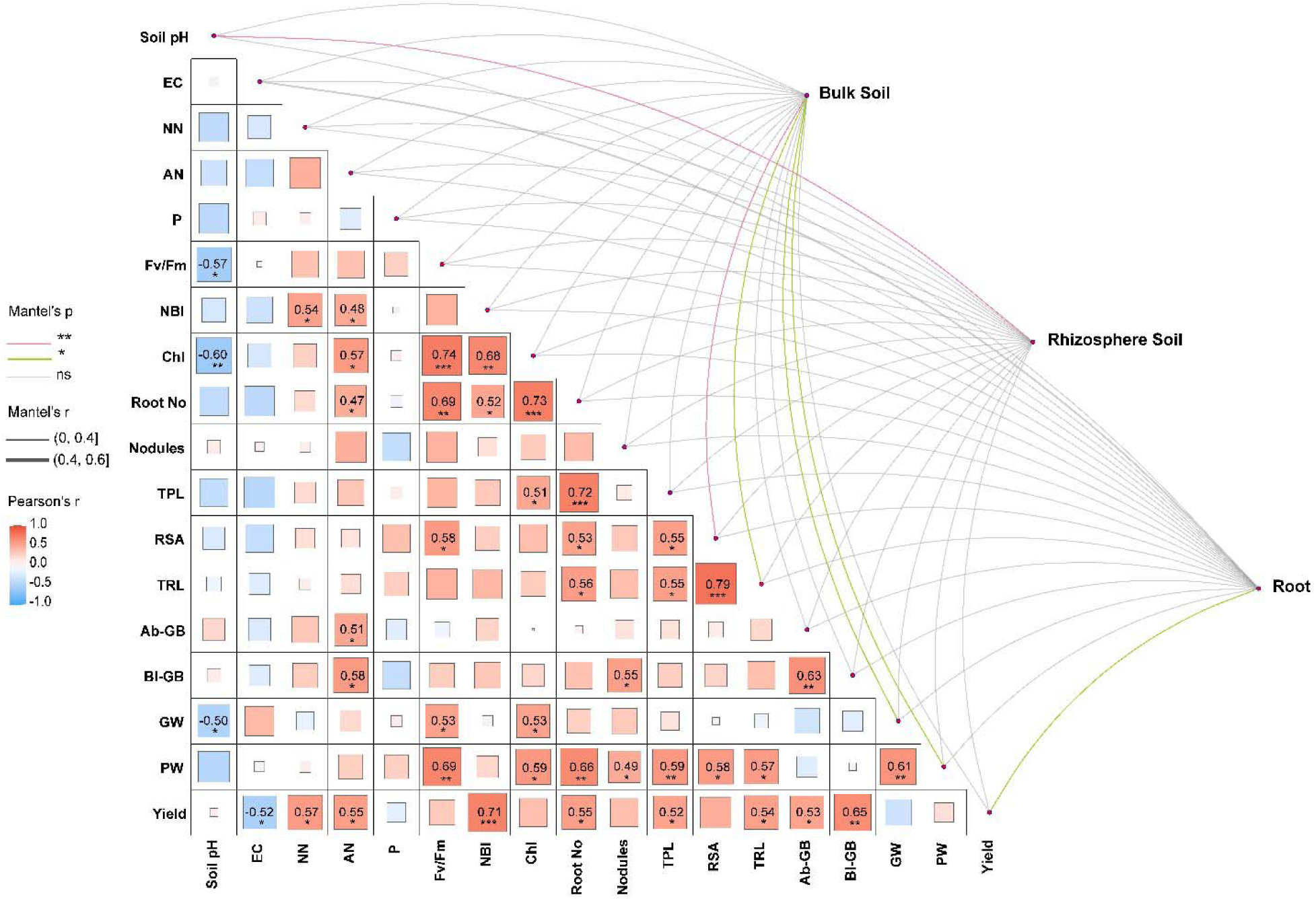
Spearman correlation and Mantel test analyses linking bacterial communities with agro-environmental traits across bulk soil, rhizosphere, and root compartments. The heatmap shows Spearman correlation coefficient (Pearson’s r) among various traits. Positive correlations are shown in red, while negative correlations are in blue, with the color gradient indicating the strength of correlation. The Mantel test analysis evaluates the strength of association between bacterial communities in each compartment. Abbreviations: EC, electrical conductivity; NN, nitrate nitrogen; AN, ammonium nitrogen; P, Phosphorus; NBI, nitrogen balance index; Chl, chlorophyll content; TPL, taproot length; RSA, root surface area; TRL, total root length; Ab-GB, aboveground biomass; Bl-GB, belowground biomass; GW, 100-grain weight; PW, 100-pod weight.

### Root bacterial networks exhibited strong association with plant traits

To gain deeper insights into the bacterial interactions across different rhizocompartments and their relationship with plant performance, co-occurrence network analysis was first conducted across root, rhizosphere, and bulk soil (**Fig. 6**). The root network displayed the highest complexity, attaining 100 nodes and 129 edges with a modularity score of 0.58, suggesting more densely connected bacterial communities. Dominant modules in the root compartment included Module #1 (16%) and Module #2 (12%), indicating closely clustered bacterial taxa. In contrast, bacterial networks in the rhizosphere and bulk soil were more fragmented and exhibited weaker interactions. Specifically, the rhizosphere network had 76 nodes and 44 edges with a modularity score of 0.945, while bulk soil had 98 nodes and 54 edges with a modularity score of 0.973. These findings highlight that root compartments harbor more structured and potentially beneficial bacterial networks than surrounding soils.

**Fig. 6.**
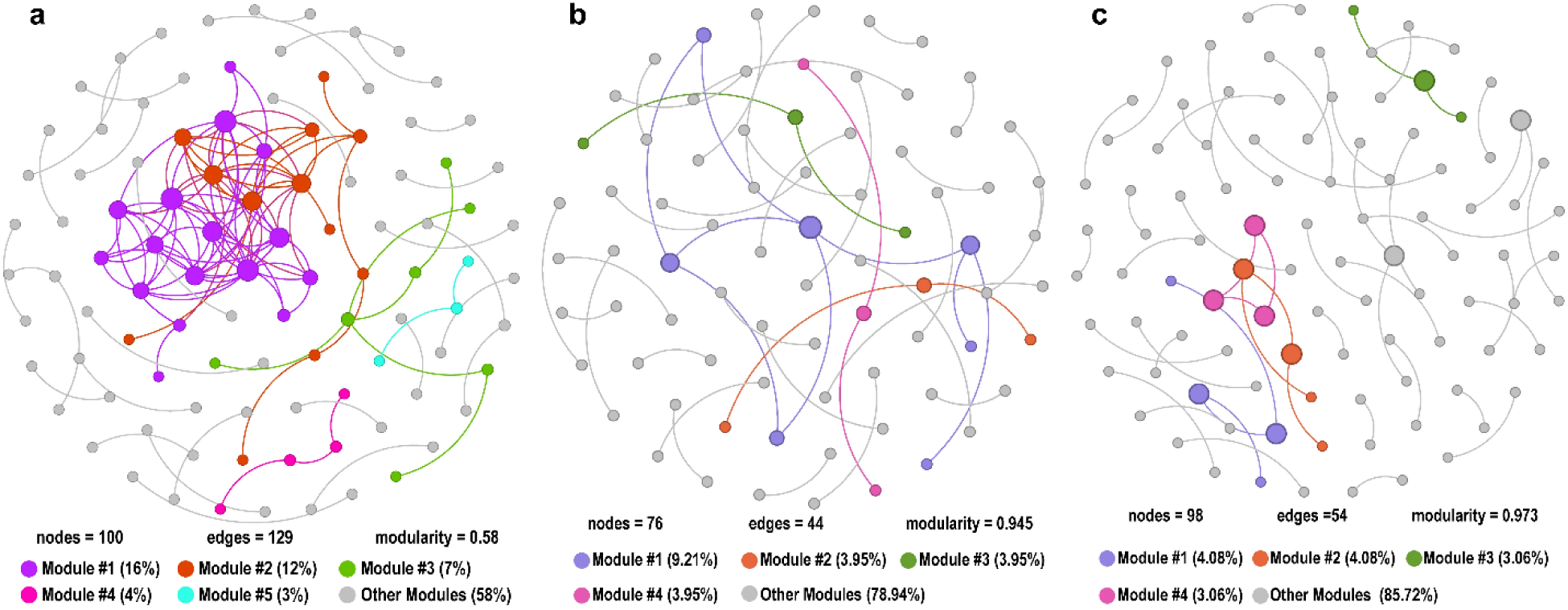
Co-occurance network analysis of bacterial communities across three rhizocompartments i.e., (a) Root, (b) Rhizosphere soil, and (c) Bulk soil. Each node represents an ASV and edges indicate significant co-occurrence associations. Colored nodes representing major modules with their respective percentages. Root compartment exhibited more complex co-occurrence network with a higher proportion of clustered modules. Gray colored nodes indicate taxa assigned to smaller or unclassified modules (Other Modules).

Considering the distinct bacterial interactions in the root compartment, we further examined the relationship between the relative abundance (Z-scores) of root modules and agro-environmental traits (**Fig. 7**). Module #1 and Module #2 were negatively correlated with most traits, whereas, Module #3 was positively associated with nitrate nitrogen, nodule number, belowground biomass, and yield. Module #5 indicated moderate correlations with a number of traits. Notably, Module #4 emerged as the most functionally relevant, exhibiting strong and significant positive correlations with majority of the traits. Taxonomic analysis revealed that Module #4 was predominantly enriched with Proteobacteria, a phylum widely recognized for its plant growth promoting and nitrogen fixing capabilities (**Table S2**). Altogether, these findings suggest that melatonin-responsive Proteobacteria in Module #4 may play pivotal role in modulating root function and plant productivity.

**Fig. 7.**
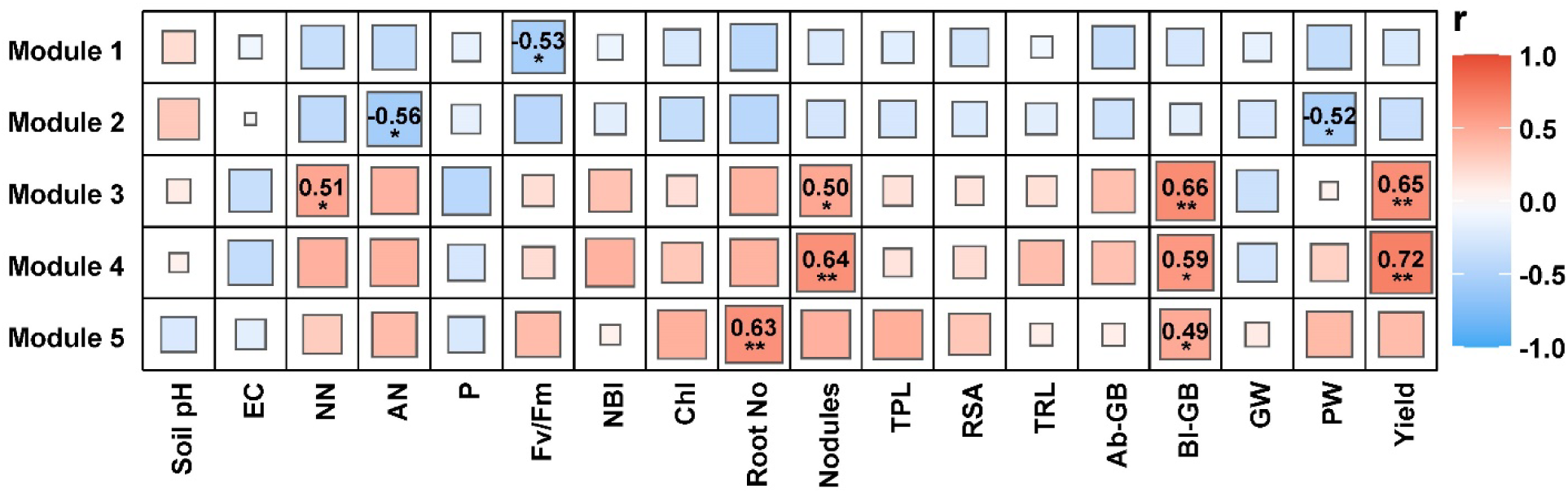
Correlation heatmap between major root modules and various agro-environmental traits. The color gradient indicates the strength (\**p* < 0.05, \*\**p* < 0.01) of correlations, where red represents positive and blue shows negative correlations. Abbreviations: EC, electrical conductivity; NN, nitrate nitrogen; AN, ammonium nitrogen; P, Phosphorus; NBI, nitrogen balance index; Chl, chlorophyll content; TPL, taproot length; RSA, root surface area; TRL, total root length; Ab-GB, aboveground biomass; Bl-GB, belowground biomass; GW, 100-grain weight; PW, 100-pod weight.

## Discussion

### Melatonin enhances root growth and yield-related traits in a genotype-specific manner

This study provides compelling evidence that exogenous melatonin enhances root development and yield-related traits in peanut, with differential responses across genotypes (Ren *et al*., 2025). Significant improvements in nodulation, root biomass, and yield related traits, particularly in KN308 and XBH16, suggest that melatonin act as a positive regulator of plant growth (**Fig. 1**). For instance, KN308 indicated the highest yield gain (151.53 to 187.04 g/m^2^), alongside enhanced root biomass and pod weight. These outcomes reveal the potential of melatonin as a bio-stimulant capable of improving agronomic performance (Wang *et al*., 2022). In soybean, melatonin application substantially increased both the number and size of root nodules by enhancing genistein accumulation in roots (Wang *et al*., 2024a). Beyond nodulation, melatonin also enhances root architecture, thereby contributing greater yield potential. In cotton, for instance, melatonin alleviated drought-induced root inhibition by promoting root length, surface area and lateral root growth, effectively mitigating yield loss (Zhu *et al*., 2023). Similarly, in Arabidopsis, melatonin enhanced primary root growth largely dependent on auxin biosynthesis and transport pathways (Yang *et al*., 2021). Moreover, melatonin-mediated yield improvement is mainly dependent on hormonal signaling including elevated levels of IAA, GA, and cytokinin while reducing ABA, to support efficient grain development (Ahmad *et al*., 2021). These findings underscore the multifaceted role of melatonin in plant growth and productivity. Functioning as a master regulator, melatonin modulates hormonal crosstalk, antioxidant defense response and stimulate nitrogen metabolism (Arnao and Hernández-Ruiz, 2019; Sun *et al*., 2021). Such mechanisms likely underline the observed improvements in chlorophyll content, nitrogen balance index, and biomass accumulation across melatonin treated plants (**Table 1**).

Importantly, the distinct response across genotypes reveals melatonin effects are modulated by intrinsic genetic factors (Wang *et al*., 2024c; Zargar *et al*., 2025). For instance, KN308 and XBH16 responded strongly, whereas XBH21 indicated a more limited improvements, suggesting potential differences in melatonin response. Such genotype-specific responses to melatonin have also been reported in other crops, influencing soil microbial abundance and activity (Li *et al*., 2025; Maleki *et al*., 2024). These findings highlight the importance of genotype-dependent strategies for melatonin application in crop improvement programs.

### Melatonin reshapes root-associated bacteria through compartment specific recruitment

In addition to its physiological effects on plant, melatonin significantly influenced the structure and composition of bacterial communities across different rhizocompartments. PCoA and PERMANOVA confirmed the distinct compartmentalization of bacterial abundance, with root-associated bacteria showing the strongest response to melatonin, particularly in XBH16 and KN308 (**Fig. 2**). This may suggest that melatonin coordinates interactions between soil microbiome and root exudates to create a friendly environment, as reported in peanut (Zheng *et al*., 2025), soybean (Xiao *et al*., 2022), apple (Cao *et al*., 2024b), and barley (Ye *et al*., 2022). Root exudate composition is known to shift in response to hormonal crosstalk, influencing the biochemical environment of the rhizosphere and thereby regulating microbial colonization (Zhalnina *et al*., 2018). Furthermore, LEfSe results revealed an enrichment of Proteobacteria related taxa, including *Rhizobium*, *Ensifer*, and *Pararhizobium* in melatonin-treated roots, contrasting with increased abundance of Firmicutes and Bacteroidota under control conditions (**Fig. 2c**). These findings are in line with emerging evidence that melatonin acts as microbiome modulator, selectively recruiting beneficial microbial taxa (He *et al*., 2023; Zheng *et al*., 2025). Specifically, Cao *et al*. (2024a) reported a melatonin-induced proliferation of Proteobacteria in the apple rhizosphere, underscoring melatonin role in the targeted recruitment of plant beneficial microbes. Gamalero and Glick (2025) further revealed that these shifts are likely driven by melatonin effects on root exudation, antioxidant response, and plant immunity, creating more favorable for beneficial taxa like Proteobacteria. Similarly, Jiang *et al*. (2022) found that melatonin favored root-microbiome interactions by regulating root development and microbial colonization, thereby contributing to better plant growth and resilience.

Plant compartment is a primary determinant in shaping the assembly of plant-associated microbiomes (Beckers *et al*., 2017; Edwards *et al*., 2015; Xiong *et al*., 2021). The root compartment indicated the highest proportion of unique ASVs (45.9%) under melatonin, with less overlap across treatments and genotypes, suggesting strong host-microbe specificity. The increased complexity of this compartment indicates enhanced bacterial interactions and cooperation, probably among beneficial taxa recruited under melatonin. A similar trend has been observed in alfalfa, where root-associated microbes respond distinctly to genotypes under stress (Fan *et al*., 2023). Complementary to previous findings that host specificity through compartment niches and genetic background plays a prominent role in shaping plant microbiome assembly (Dicenzo *et al*., 2016; Gao *et al*., 2020), our study provides new evidence that plant compartments significantly influence not only microbiome assembly but also their functions. Genotype-specific microbial assemblage has also been reported in barley and wild perennial plants, where differences in host genotype substantially influences microbial colonization and function (Bulgarelli *et al*., 2015; Wagner *et al*., 2016). Moreover, compartments-specific changes have been observed in barley, where melatonin application significantly reconstructed microbial community structure, altering the abundance of major bacterial taxa (Ye *et al*., 2022). Such compartmentalization and genotype-specific microbial recruitment may therefore be vital to the functional reshaping of root microbes under melatonin treatment, with implications for nutrient acquisition, growth promotion and stress resilience (Compant *et al*., 2019; Toju *et al*., 2018).

### Root bacterial networks link melatonin enriched taxa to enhanced plant productivity

The plant and its associated microbiome function as a meta-organism, often referred to as the holobiont (Berg *et al*., 2021). The dynamic assemblage, comprising millions of microbial inhabitants, forming a complex community that significantly influences plant productivity through collective metabolic activities and host interactions (Schmidt *et al*., 2014). The network analysis of co-occurrence taxonomic patterns provides valuable insights into the assembly mechanisms of bacterial communities (Chen *et al*., 2018). In our study, co-occurrence network analysis highlighted that melatonin application not only affect community composition but also enhance the structural connectivity of root-associated bacterial networks. The root compartment displayed the most densely connected network (100 nodes, 129 edges), with several well-defined modules (**Fig. 6a**). The increased number of nodes and edges in this compartment suggest mores interconnected bacterial communities (Kajihara and Hynson, 2024). Module #4 emerged as functionally significant, positively correlating with traits such as nitrate nitrogen, nodulation, root biomass, and yield. This module was enriched with Proteobacteria, highlighting their role as key microbial taxa in melatonin-enhanced plant performance (**Table S2**). The functional significance of this module is further supported by previous literature underscoring Proteobacteria as key microbial players in plant performance (Mendes *et al*., 2013). For instance, Yaghoubi Khanghahi *et al*. (2021) isolated *Pseudomonas*, *Comamonas*, and Acinetobacter, all belonging to Proteobacteria, indicating strong growth-promoting traits including nitrogen fixation, directly aligned with the plant traits associated with module #4 in our study. Similarly, Di Benedetto *et al*. (2017) reported the significance of *Rhizobium* and other Proteobacteria in regulating nitrogen use efficiency in wheat, emphasizing their role in sustainable plant-microbe interactions. The integration of these bacterial taxa into densely clustered co-occurrence modules indicates that melatonin may not only recruit these beneficial bacteria but also improve ecological conditions that favor their cooperative crosstalk.

Highly modular and interactive networks, like those detected in root compartment, are often associated with functional redundancy, ecological stability, and enhanced stress resilience (Luo *et al*., 2021; Toju *et al*., 2018). Moreover, the tighter and more consistent structure of the root bacterial network under melatonin implies enhanced bacterial cooperation or co-dependency, potentially facilitating beneficial functions such nitrogen fixation, hormone biosynthesis, and growth promotion. Specifically, the enrichment of *Sphingomonas* and *Enterobacter hormaechei*, recognized for their plant-growth promoting and biocontrol capabilities, in melatonin treated roots (**Fig. S2**) further support this interpretation (Ma *et al*., 2016). *Sphingomonas* has been reported to enhance disease resistance through symbiotic interactions triggered by root exuded succinic acid, thereby regulating nutrient cycling and microbial network complexity (Shi *et al*., 2022; Wang *et al*., 2025). Similarly, *E. hamaechie* contributes to root development through nitrogen fixation, phosphate and potassium solubilization, while also participating in disease suppression by reshaping root-associated microbes (Ranawat *et al*., 2021; Wang *et al*., 2024b). Notably, the rhizosphere and bulk soil compartment remained sparse and revealed weaker trait associations, emphasizing the importance of root compartment where melatonin exerts its microbiome-mediated benefits. Together, these findings suggest that melatonin enriched functionally active and densely connected bacterial taxa, in which Proteobacteria form the central nodes linked to key physiological traits.

### Conclusion

This study provides comprehensive evidence that exogenous melatonin application enhances peanut productivity and root associated microbiome dynamics in a genotype and compartment specific manner. Melatonin not only improved physiological and agronomic traits such as nodulation, root biomass, and yield associated traits, but also significantly altered the composition of bacterial communities, particularly with the root compartment. Responsive genotype like Kainong 308 indicated distinct improvements in root biomass and yield gain, highlighting melatonin potential as bio-stimulant in legume crops. Microbiome analysis revealed that melatonin selectively enriched beneficial taxa, especially Proteobacteria such as *Rhizobium*, *Sphingomonas*, and *Enterobacter hormaechei*, with in the root compartment. These bacteria formed densely connected co-occurrence modules, with module #4 strongly correlating with most of the plant traits. Together, these findings reveal melatonin functions as both plant growth regulator and microbiome modulator. This dual role underscores melatonin potential in sustainable agriculture through coordinated improvements in plant performance and microbiome-mediated resilience. Future studies should explore how melatonin-mediated shifts in root exudates shape microbiome recruitment under stress conditions. Moreover, microbiome engineering strategies targeting melatonin-enriched taxa hold promise for crop improvement across diverse environments.

## Supplementary data

The following supplementary data are available at JXB online.

**Fig. S1.** Rarefraction curve. X-axis indicates counts of randomly sampled sequences, while Y-axis shows counts of features detected by given sequences. Lines with different colors stand for different smaples given

**Fig. S2.** LEfSe cladogram showing differentially abundant bacterial taxa across treatments, genotypes, and rhizocompartments. Cladogram illustrates the taxonomic arrangement from phylum to genus level, highlighting bacterial lineages significantly enriched in specific sample groups. Each colored node represents a taxon significantly associated with a particular genotype (Xinbaihua 16, Xinbaihua 21, or Kainong 308), rhizocompartment (bulk soil, rhizosphere, and root), and treatment (control or melatonin). These results suggest a strong genotype- and compartment-specific influence of melatonin on microbial abundance.

**Fig. S3.** Taxonomic composition of bacterial communities in peanut rhizocompartments at (a) Class, (b) Order, (c) Family, and (d) Genus levels under control and melatonin treatments. Relative abundance profiles of bacterial communities are shown for root (R), bulk soil (BS), and rhizosphere soil (RS) compartments of three peanut varieties, Xinbaihua 16, Xinbaihua 21, and Kainong 308 under control (CK) and melatonin (MT) treatments. Taxa with low abundance were grouped under “Others” or “Unassigned”. The plots highlight compositional shifts in the microbial community structure across genotypes, compartments, and treatments.

**Table S1.** Relative abundance of major bacterial phyla in roots, rhizosphere, and bulk soil across genotypes under control (CK) and melatonin (MT) treatments.

**Table S2.** Taxonomic composition of dominant bacterial phyla within major root co-occurrence network modules

## Author Contributions

AM: conceptualization, investigation, writing - original draft; XK: methodology, formal analysis; LL: data curation; MHUK: visualization; PJ: resources; CM: writing - review and editing; ZZ: supervision, funding acquisition.

